# Rapid responses of winter aphid-parasitoid communities to climate warming

**DOI:** 10.1101/411975

**Authors:** K. Tougeron, M. Damien, C. Le Lann, J. Brodeur, J. van Baaren

## Abstract

Consequences of environmental fluctuations, including those associated with climate changes, can have a knock-on effect from individual to community scale. In particular, changes in species seasonal phenology can modify the structure and composition of communities, with potential consequences on their functioning and the provision of ecosystem services. In mild climate areas, aphids can be present in cereal fields throughout the winter, which allows aphid parasitoids to remain active. Using a nine-year dataset of aphid-parasitoid winter trophic webs in cereal fields of Western France, we report that the community structure and composition that prevailed before 2011 have recently shifted toward a more diversified community, with the presence of two new braconid parasitoid species (*Aphidius ervi* and *Aphidius avenae*), a few hyperparasitoid species and one aphid species (*Metopolophium dirhodum*). Increases in minimal winter temperatures and the frequency of frost events across the years partially explain observed community changes. Strong bottom-up effects from the relative abundances of aphid species also determine the relative abundance of parasitoid species each winter. Strong compartmentalization in parasitoid preference for host is reported. We suggest the recent modifications in parasitoid community composition to be linked to shifts in diapause expression (reduction or arrest of the use of winter diapause) and to host availability throughout the year. We highlight the implications for natural biological control in cereal fields. Perspectives are proposed to predict the composition of future host-parasitoid communities in the climate change context.

## Introduction

Climate change impacts the geographic distribution, diversity and abundance of organisms (Walther et al., 2002; Parmesan, 2006). In particular, climate warming can strongly influence their seasonal phenology, migration pattern, number of generations per year and overwintering strategy (Roy and Sparks, 2000; Altermatt, 2010; Bale and Hayward, 2010). In temperate areas temperatures are increasing faster in winter than in summer, leading to overall milder, shorter and later winter periods (IPCC, 2014). Plastic and adaptive responses of organisms to new thermal environments could modify species interactions such as competition, predation and parasitism and impact the structure and stability of communities (Hughes, 2000).

In the context of the global diversity crisis, studies increasingly focus on how trophic networks respond to global changes (Chaianunporn and Hovestadt, 2015; Parmesan, 2006). Indeed, species interactions within and between communities support the majority of ecosystem services and must be considered as study systems *per se* (Montoya et al., 2003). In some cases, food web structure and composition are quite fragile and are likely to rapidly change in the context of climate warming; understanding how and why these food webs vary in space and time is a central objective in community ecology (Facey et al., 2014). New species appear while others disappear from the food web and changes in species interactions between trophic levels occur (Chaianunporn and Hovestadt, 2015; Tylianakis et al., 2008).

Parasites are omnipresent in almost every food web (Dobson et al., 2008) and their interactions with hosts greatly contribute to ecosystem functioning (Lafferty et al., 2008). Their ecology is tightly associated with their hosts’ and likely to be influenced by recent climate change, threatening the provisioning of ecosystem services such as natural biological pest control by insect parasitoids (Hance et al., 2007; Jeffs and Lewis, 2013). The impacts of climate change or inter-annual variations in climatic conditions on host-parasitoid communities remain little explored compared to other types of food webs (e.g. plant-herbivore networks; Singer and Parmesan, 2010) and there have been few attempts at predicting their future structure and composition under different scenarios of climate change (Jeffs and Lewis, 2013).

In regions characterized by mild winter temperatures, the absence of lethal frosts allows aphids and their parasitoids to remain active and reproduce throughout winter. In cereal crops of Western France, aphid-parasitoid communities in winter were historically composed of the two parasitoid species *Aphidius rhopalosiphi* De Stefani-Perez and *Aphidius matricariae* Haliday and the two aphid species *Rhopalosiphum padi* (L.) and *Sitobion avenae* (Fabricius). From late spring to fall, additional species were present, including the parasitoids *Aphidius ervi* Haliday and *Aphidius avenae* Haliday (Krespi, 1990; Krespi et al., 1997; Rabasse et al., 1983) and the aphid *Metopolophium dirhodum* (Walker). These seasonal variations in aphid and parasitoid species occurrence seem to be consistent across Western Europe in cereal crops (Honek et al., 2018; Lumbierres et al., 2007) and reflect thermal niche separation (Andrade et al., 2016; Le Lann et al., 2011). *Aphidius avenae* parasitoid shows less cold resistance and more heat resistance than *A. rhopalosiphi* (Le Lann et al., 2011), while the aphid *R. padi* prefers cooler conditions and is more cold resistant than *S. avenae* (Alford et al., 2016; Jarošík et al., 2003; van Baaren et al., 2010).

Andrade et al. (2016) reported that in host-parasitoid communities of Western France parasitoid species usually not encountered during winter are now being active throughout the season and exploiting anholocyclic aphids (i.e., aphids having parthenogenetic reproduction all-year-long). In the present study, we have adopted a community-wide approach and analyze the effects of long-term (inter-annual) variations in climatic conditions on aphid and parasitoid species occurrence and relative abundance. Using a nine-year dataset, we explored community assembly rules by first describing temporal changes in winter aphid-parasitoid associations, and then linking these changes to modifications in winter climatic conditions and to other abiotic and biotic factors such as shifts in species interactions. We expect warmer winters to be associated with occurrence and higher abundance of *A. avenae* and *A. ervi* for parasitoids and *S. avenae* and *M. dirhodum* for aphids. We expect colder winters to be associated with the presence of *A. rhopalosiphi* and *A. matricariae* for parasitoids and *R. padi* for aphids.

## Material and Methods

### Data collection

Data consist in aphid-parasitoid pairs of species gathered from different studies conducted in the Long Term Ecological Research (LTER) ZA Armorique, France (48.29 °N – 1.35 °W), each winter from 2009/10 to 2017/18, at variable dates from late-November to mid-March of each year (excepted in 2009/10 when sampling was conducted only in January and February). Data from winter 2009/10 to winter 2012/13 were obtained from Andrade et al., (2016) and Eoche-Bosy et al., (2016), data of 2013/14 from Tougeron et al., (2016), data of 2014/15 from Tougeron et al., (2017), data of 2015/16 from Damien et al., (2017) and data of 2016/17 and 2017/18 from unpublished field results. In winter 2010/2011, no parasitoids nor aphids were found in the fields due to frost conditions during 15 consecutive days in November (Andrade et al., 2016) so this winter was excluded from the dataset to minimize unbalanced analyses on community data.

Mean, mean minimum and mean maximum daily temperature data per sampling year in the LTER were obtained from Météo France (2018). Additionally, we calculated the number of frost events (i.e., occurrence of at least three consecutive days with negative mean temperatures (which could be lethal for most species) and mean duration of frost events (days). Highly correlated variables >70% were not used for our analyses. We thus only used the mean minimal temperature (correlated with mean temperature and mean maximum temperatures) and the mean duration of frost events (correlated with the number of frost events).

### Sampling and quantitative food-webs

In each of the studies from which data were collected, sampling was performed following the protocol of Andrade et al. (2016), with differences in the location of sampled fields due to crop rotations. In brief, sampling was conducted every ten days in six to fifteen cereal fields each year; mainly winter wheat, but also barley and triticale. All aphids and aphid mummies (i.e., exoskeleton of dead aphid containing a developing parasitoid) were randomly collected during a one-hour period over an approximate area of 1000m^2^. Aphid density was very low in winter; around one aphid/m^2^. Accordingly, parasitism rate during all winters was high (60-90%), underlying rarity of hosts and high competition levels among parasitoids. As the aphid-parasitoid network is stable over the winter sampling period (Andrade et al., 2016; Damien et al., 2017), data were pooled for the entire winter season. Live aphids were brought back to the laboratory and reared on winter wheat until mummification or death. All mummies were maintained in gelatin capsules at ambient temperature (17-20°C) until parasitoid emergence. Emerging adult primary parasitoids and aphid hosts were then identified to the species based on morphological characters (Hullé et al., 2006). Hyperparasitoids (secondary parasitoids) were identified to the genera level. Parasitoids and hyperparasitoids that emerged more than 25 days after sampling, representing each year less than 25% of the total number of sampled mummies, were excluded from the food-web analysis to avoid accounting for diapausing individuals when characterizing winter-active communities. Important differences in sample sizes are due to differences in both sampling effort among years and climatic conditions.

To examine trophic interactions between host and parasitoid species, quantitative food webs using the relative abundance (%) of each species were constructed for each winter following the methodology of Memmott et al., (1994). Hyperparasitoids were assigned to the same trophic level than primary parasitoids because it was impossible to assess in which parasitoid host species they developed.

### Analyses

Food webs were compared among years using different quantitative and qualitative metrics calculated using the *bipartite* (Dormann et al., 2009) and the *codyn* R packages (Hallett et al., 2016): Connectance - the overall complexity of the food web (realized proportion of potential links); Web Asymmetry - the balance between numbers of parasitoid and aphid species (negative values indicate more species in higher than in lower trophic-level); H2 - the level of specialization within a network, from 0 (no specialization) to 1 (perfect specialization); Generality - the weighted mean number of aphid species exploited by each parasitoid species; Vulnerability - the weighted mean number of parasitoid species attacking a given aphid species.

Then, we performed a non-metric multidimensional scaling (NMDS) analysis to group years by climatic similarities based on a distance matrix; accordingly, years were groups by four based on their distance to the more extreme years on the NMDS. Following these analyses, years were characterized as either mild or cold winters, and these categories were used to group years on the following Principal Component Analysis (PCA). A first PCA was performed to separate each year of sampling based on selected climatic variables; mean minimal temperatures and mean duration of frost events. Another PCA analysis was performed to separate each year of sampling based on aphid and parasitoid species relative abundances. Finally, a canonical correspondence analysis (CCA) was performed to assess relationships between the aphid and parasitoid species and the climatic variables matrixes. ANOVA-like permutation tests for CCA (*vegan*) were used to assess the significance of constraints. Only primary parasitoids from the *Aphidius* genus were considered for analyses on climatic variations.

As we wanted to account for species co-occurrences in our analyses, species-by-species models were not appropriate. Instead, we used a community approach by analyzing separately the effects of the selected climatic variables across years on parasitoid species (matrix containing the relative abundance of the four parasitoid species) and aphid species (matrix containing the relative abundance of the three aphid species). To do so, Bray-Curtis dissimilarity indexes in species relative abundances of each sampling year were calculated (separately for parasitoids and aphids) and fitted to linear models as explanatory variables using the Adonis-Permanova function from the R package *vegan* (Oksanen et al., 2015) calculating permutation test with pseudo-F ratios. For aphids, the minimal temperature and the mean duration of frost events were used as explanatory variables. For parasitoids, we also included the relative abundance of each of the three aphid species as explanatory factors in the linear models. All analyses were performed using R software (R Core Team, 2017).

## Results

Changes in species richness and relative abundances were observed from winter 2009/10 to winter 2017/18 in the aphid-parasitoid food webs, with important inter-annual variations (Fig. 1A). The year 2009/10 was similar to the past three decades, as described in introduction, with *A. rhopalosiphi* and *A. matricariae* being the only two parasitoid species active in winter and exploiting *S. avenae* and *R. padi*. In addition to these two aphid species, the aphid *M. dirhodum* was present every winter starting 2011/12 at 12%, and it represented up to 68% of the aphid species relative abundance in winter 2015/16 (Fig. 1A). There was high variability in aphid proportions between years, for each species.

**Figure 1:**
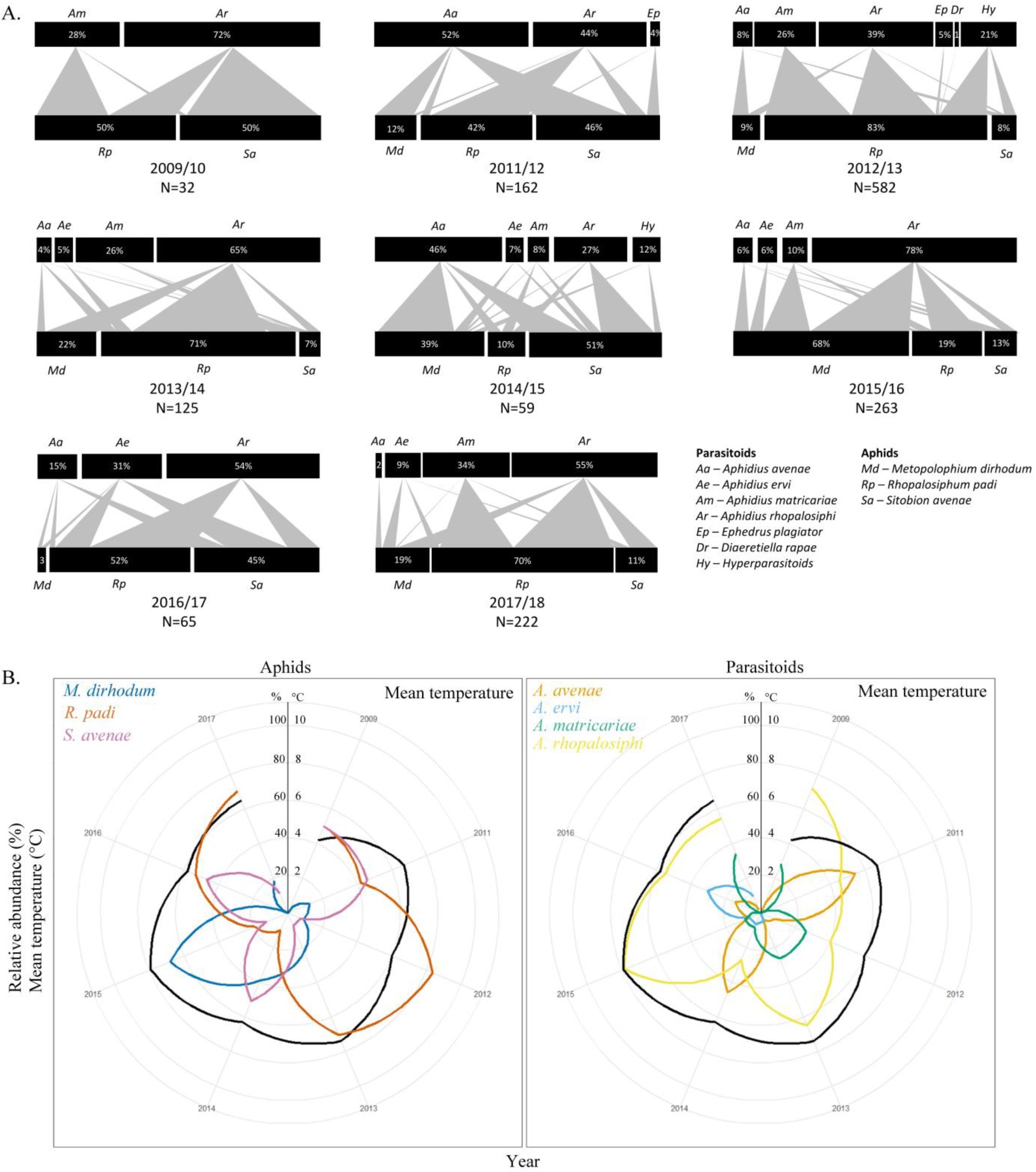
**A.** Quantitative food webs of parasitoid and aphid community composition in winters 2009/10 to 2017/18 (there was no insects in 2010/11). Upper and lower bars represent the relative abundance (%) of parasitoid, including hyperparasitoids, and aphid species, respectively. The thickness of the arrow between bars is proportional to the interaction strength between a pair of species. Total number of individuals (N) used to construct each food web is shown for each year. **B.** Rankplots showing the relative abundances (%) of aphids (left panel) and *Aphidius* parasitoids (right panel) each winter from 2009/10 to 2017/18. Mean winter temperature (°C) is shown in black.

*Aphidius avenae* was observed for the first time in the winter 2011/12 with a relative abundance of 52%. *Aphidius ervi* was observed in the network in 2013/14 with a relative abundance of 5%. Both species have since remained present in the network, although at variable relative abundances. *Aphidius rhopalosiphi* was present every winter while *A. matricariae* occurrence and relative abundance were highly variable over the years. *Ephedrus plagiator* (Nees) and *Diaeretiella rapae* (M’intosh), two generalist species (Hullé et al., 2006), were anecdotally reported in winter 2011/12 and 2012/13. Hyperparasitoids from the genera *Alloxysta*, *Asaphes* and *Phaenoghyphis* were also detected in two out of eight winters.

Mean temperature varied across years and ranged from 4.2°C in 2009/10, to 7.9°C in 2015/16, and was associated with variations in aphid and parasitoid relative abundances (Fig. 1B).

Food-web metrics are summarized in Table 1. The food-web connectance was ≥0.6 every year; all links the first year and almost all potential links the following years between each parasitoid and aphid species were observed. In most years, the food web was asymmetric, with more parasitoid species than aphid species. The degree of specialization within the food-web (H2 index) tended to decrease over the years. Accordingly, the generality and vulnerability indexes tend to increase over time; each parasitoid species attacked more aphids and each aphid species was exploited by more parasitoid species over the years, in general (Table 1).

**Table 1:**
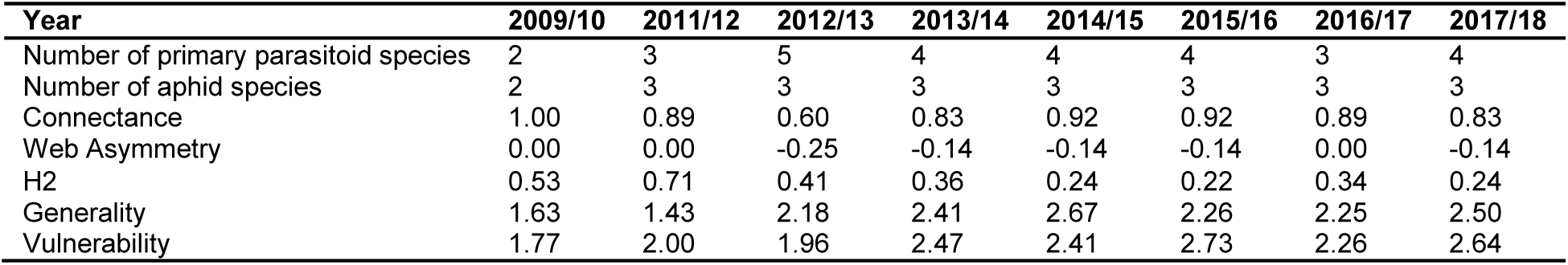
Number of species in each trophic level and food-web metrics for each sampled winter.

NMDS analysis showed that years 2009/10, 2012/13, 2013/14 and 2017/18 can be grouped together as “cold winters” whereas years 2011/12, 2014/15, 2015/16 and 2016/17 can be described as “mild winters”. This clustering was supported by 94.7% inertia on the PC1 of the PCA analysis (Fig. 2A). Graphically, coldest winters (i.e. decreasing minimal temperatures and increasing duration of frost events) were overall associated with the co-occurrence and higher relative abundances of both *A. rhopalosiphi* and *A. matricariae* parasitoids and *R. padi* aphids, while warmest winters were associated with co-occurrence and higher abundance of both *A. avenae* and *A. ervi* parasitoids and *S. avenae* aphids. *Metopolophium dirhodum* was highly abundant in the warmest winter (2015/16) (Figs. 1, 2B). However, the aphid-parasitoid community PCA was only supported by 43% inertia on the PC1, indicating that species partition on this figure is only partially explained by the sampling year (Fig. 2B).

**Figure 2:**
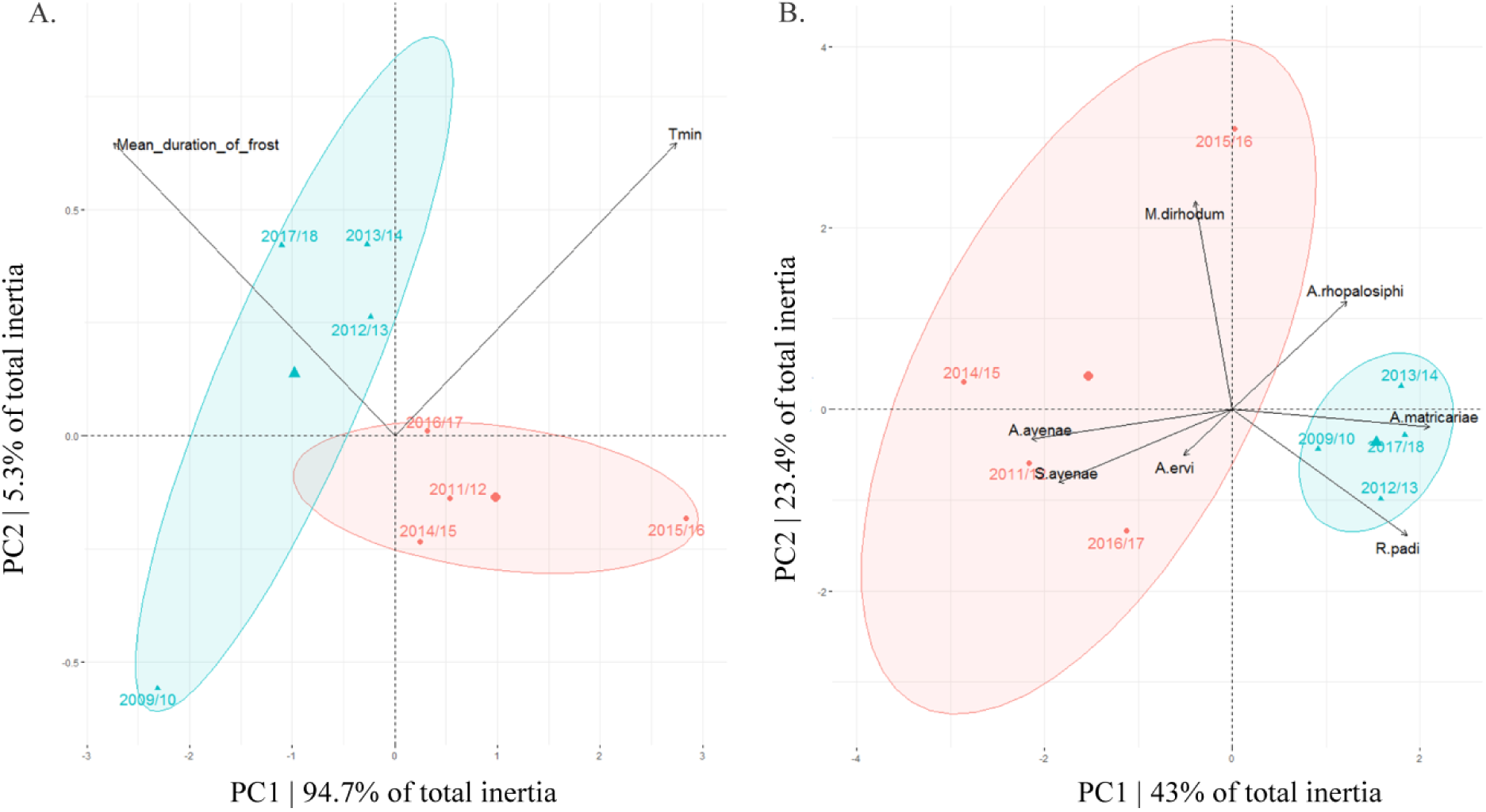
**A.** PCA partitioning two climatic variables (mean minimal temperatures “Tmin” and mean duration of frost events) by sampling years. **B.** PCA partitioning aphid and parasitoid species relative abundances by sampling years. Confidence ellipses are constructed around groups of cold (blue) and mild (red) winters.

We found a marginally non-significant influence of the selected climatic data (mean minimal temperature and duration of frost events) on global aphid and parasitoid relative abundances across years (CCA ANOVA-like permutation test, F=1.67, df=2, p=0.06). In details, aphid abundances were significantly affected by changes in mean minimal temperatures (Permanova, F=2.03, df=1, R2=0.22, p=0.03) and in mean duration of frost events across the years (F=2.57, df=1, R2=0.27, p=0.04), in the way described in the precedent paragraph. Parasitoid abundances were marginally significantly affected by changes in mean minimal temperatures (F=2.67, df=1, R2=0.20, p=0.05) and significantly affected by changes in mean duration of frosts across the years (F=14.72, df=1, R2=0.15, p=0.01), as described above. Changes in parasitoid abundances across the years were not affected by the abundances of the aphid *S. avenae* (F=1.7, df=1, R2=0.16, p=0.16) but was significantly affected by the abundances of *M. dirhodum* (F=35.5, df=1, R2=0.37, p=0.01) and *R. padi* (F=17.2, df=1, R2=0.18, p=0.009).

Preferential associations between aphid hosts and primary parasitoids species occurred as shown by the arrows on Fig. 2B (pooled data across years). The aphid *S. avenae* was mostly associated with the parasitoids *A. avenae* and *A. ervi* while the aphid *R. padi* was mostly associated with the parasitoids *A. rhopalosiphi* and *A. matricariae*. *Metopolophium dirhodum* was not preferentially associated with any parasitoid in the food web (Fig. 2B).

## Discussion

Our results illustrate how climatic changes during winter have rapidly, over nine years, translated into modifications in species composition within an aphid-parasitoid community. Two parasitoid species *A. avenae* and to a minor extent *A. ervi,* and one aphid species *M. dirhodum* are now active in Western France throughout winter together with other species of the community. The winter trophic network composition in cereal fields is getting similar to what is usually described in spring in this area. The winter food web has become more diversified in aphid and parasitoid species and, while the connectance (realized links) remains stable over time, the degree of specialization tends to decrease, suggesting that parasitoids exploit aphids in function of their relative abundance, as reported in spring (Andrade et al., 2016). This may be due to increasing aphid densities in winter on cereal crops, leading to lower competition pressure among parasitoids.

Changes in occurrence do not arise from recent modifications in distribution range of the species, since *A. avenae*, *A. ervi* and *M. dirhodum* have been commonly observed for several decades in spring at the same location (Andrade et al., 2016; Krespi, 1990). Our results suggest a recent shift in overwintering strategy in *A. avenae* and *A. ervi* parasitoid populations with some individuals remaining active throughout the winter rather than entering diapause. This hypothesis is supported by results from a laboratory experiment showing that diapause incidence in both parasitoid species was low (<15%), even when parasitoids were reared under fall-like temperature conditions that usually induce high levels of diapause (Tougeron et al., 2017). Variations in species composition in the food web over the years may arise from differences in thermal niches; the most cold-resistant species usually remained active during winter (e.g., *A. rhopalosiphi*, *A. matricariae* and *R. padi*) whereas less cold-resistant species (e.g., *A. avenae*, *A. ervi*, *M. dirhodum* and hyperparasitoids) were probably in diapause mostly active from spring to fall (Alford et al., 2016; Andrade et al., 2016; Krespi, 1990; Le Lann et al., 2011; Tougeron et al., 2017). Overwintering temperature may now be warm enough to allow niche overlapping of all these species during winter.

It has been shown that fine-scale intra-seasonal temperature variations (i.e. temperature experienced by the insect during its development) played an important role in shaping local aphid-parasitoid communities in Western France (Andrade et al., 2015, 2016). For example higher developmental temperatures were associated with increasing abundance in *A. avenae, S. avenae* and decreasing abundance in *R. padi* (Andrade et al., 2016). Winter 2016/17 was on average warmer than other winters but important cold spells occurred in December and January, which may have conducted to higher abundances of *R. padi* and *A. rhopalosiphi*, the more cold tolerant species in the food-web (Alford et al., 2016; Le Lann et al., 2011), and quasi-disappearance of *M. dirhodum* from the system, through environmental filtering. Such thermal extremes and microclimatic variations may reduce or eliminate any advantages of global warming for some species (Ma et al., 2015; Sgrò et al., 2016) and may impede evaluation and prediction of climate change effects on community dynamics (Bailey and van de Pol, 2016; Blonder et al., 2017).

We have shown that both mean minimal temperatures and mean duration of frost events over the winter are predictors of winter aphid abundances and of their variation in occurrence among years. Honek et al. (2018) also demonstrated that temperature in winter was an important predictor of maximum abundances of cereal aphids during the weeks following sampling. However, change in mean minimal temperature and duration of frost events only slightly contributed to the trend observed in parasitoid relative abundance changes over the years. Stochastic effects or other environmental variables than temperature such as host-parasitoid interactions may better explain inter-annual variations in species abundances and occurrences during winter. For instance, we showed high level of host-parasitoid compartmentalization within the food web; the variation in relative abundance of some species was highly correlated with abundance of other species, suggesting bottom-up effects on parasitoid abundance. The importance of host species in shaping parasitoid response to climate warming must therefore be accounted (Barton and Ives, 2014). Parasitoids and their hosts may also be influenced by microclimatic refuges in the landscape (Alford et al., 2017; Tougeron et al., 2016), by surrounding plant covers (Damien et al., 2017; Gagic et al., 2012) or by plant quality (Honek et al., 2018).

Modifications of the parasitoid guild could also be due to shifts in competition for hosts following the addition of new species. Indeed, female parasitoids show seasonal variations in foraging behavior (Roitberg et al., 1992) and can adapt their foraging strategies to competition or host-patch quality (Barrette et al., 2010; Le Lann et al., 2008; Moiroux et al., 2015; Outreman et al., 2005). In winter, it has been demonstrated that female parasitoids adopt generalist strategies due to shortage of optimal hosts, leading to high competition, whereas spring parasitoids usually display specialist strategies by selecting optimal host species (Eoche-Bosy et al., 2016). The recent addition of *A. avenae* and *A. ervi* in the overwintering food web, which are good competitors at exploiting *S. avenae*, may have reduced the abundance of *A. matricariae* and *A. rhopalosiphi* (Andrade et al., 2016; Eoche-Bosy et al., 2016; Le Lann et al., 2012).

Climate warming challenges the coexistence and interactions between ecologically related species, as well as community stability and ecosystem functioning (Tylianakis et al., 2008; van der Putten et al., 2004). In cereal fields, overwintering reproduction in aphid parasitoids plays an important role in suppressing early cereal aphid populations in spring (Honek et al., 2018; Langer et al., 1997; Plantegenest et al., 2001). Increasing number of parasitoid species during winter due to climate warming could enhance aphid natural biological control through increasing already high parasitism rate in winter, even if consequences of niche overlapping between parasitoid species via addition of new species in the food-web are difficult to predict. The presence of non-diapausing hyperparasitoids, reported for the first time in winter in Western France in 2012/13 (Tougeron et al., 2017), may reduce the efficiency of biological control in the fields, although hyperparasitoids can sometimes stabilize primary parasitoid populations (Tougeron and Tena, 2018). In Spain, characterized by relatively warm winters, hyperparasitism remains high throughout the year which disrupts biological control by primary parasitoids in orchards (Gómez-Marco et al., 2015). With an expected temperature increase from 0.5 to 2 °C in the next decades (Karl and Trenberth, 2003), occurrences of new species in food-webs such as hyperparasitoids can be more common, either through shifts in geographic distribution (e.g. biological invasions) or shifts in phenology (e.g. reduction of diapause expression) (Tougeron and Tena, 2018).

Based on the data currently available on this host-parasitoid system, we observed the homogenization of the winter and spring aphid-parasitoid communities, mostly due to change in phenology after increasing temperatures during winter and decreasing duration of frost events. High variations between years may underline a transition period between two episodes of stable communities, although the food-web could also remain unstable due to variations in climate at fine temporal scale. Predictive analyses on the community structures should now integrate local changes in overwintering strategies of one or more species to identify the potential effect of climate change on the ecosystem service provided by parasitoids.

## Acknowledgments

We thank all collaborators at Ecobio lab who helped with field work, S. Chollet and J.-S. Pierre for their assistance in analyzing the community data. KT was funded by the French Region Bretagne and the Canada Research Chair in biological control awarded to JB. MD was funded by the FLEUR project and INRA. This study was supported by the LTER France Zone Atelier Armorique. KT and MD collected and analyzed the data. KT wrote a first version of the manuscript. All coauthors made substantial contributions to the manuscript.

